# Dietary sterol depletion is associated with gut dysfunction in *Drosophila melanogaster* females

**DOI:** 10.1101/2025.02.11.637748

**Authors:** Brooke Zanco, Christen K Mirth, Carla M Sgro, Matthew DW Piper

## Abstract

Recent findings have shown that dietary restriction (DR) may extend lifespan by improving late life gut health. Given that recent work has shown that micronutrients such as sterols can mediate the effects of macronutrient ratios on longevity, we decided to examine the effects of sterol limitation on gut health. We found that mothers fed low sterol diets both suffer from increased incidence of intestinal permeability and reduced lifespans. Moreover, epithelial cells in the anterior region of the guts of flies fed low sterol diets showed signs of dysfunction prematurely when compared to flies fed adequate sterols. These findings indicate that the relationship between gut dysfunction and death in *D. melanogaster* females is conditional on dietary sterol limitation and may point to different paths of physiological decline with ageing that depend on the precise nutritional composition of the diet.

## Introduction

Diet influences many fitness-related traits, including lifespan and fecundity (Partridge, Gems and Withers, 2005; Barnes *et al*., 2008). Specifically, dietary restriction (DR), whereby food intake is reduced without malnutrition, has been shown to extend lifespan across a broad range of taxa (McCay *et al*., 1935; Le Couteur *et al*., 2014). Because of this, dietary interventions are thought to be one of the most promising means of extending lifespan. Nevertheless, the mechanistic basis through which DR extends lifespan remains elusive.

The physiological changes that occur in different organs throughout an organism’s lifespan is a fundamental question in ageing research, with a mounting body of work aiming to understand the ways in which the health of specific organs mediates lifespan outcomes (Garigan et al., 2002; LA et al., 2002; Wessells et al., 2004; Biteau et al., 2010). Recent work suggests that physiological decline in specific organs is causative of death, and that the preservation of these organs can in turn, prolong lifespan (Garigan *et al*., 2002; LA *et al*., 2002; Wessells *et al*., 2004; Biteau *et al*., 2010; McGee *et al*., 2011). One organ thought to be central in mediating lifespan in *Drosophila melanogaster* and other organisms is the gut, whose epithelial barrier function declines with age causing a leaky gut phenotype (Salazar *et al*., 2023). These effects have been predominantly reserved to females in *Drosophila* (Regan *et al*., 2016), which is thought to be caused by female specific gut remodelling in response to mating and diet, which are the same conditions that modify lifespan. In contrast, males show little gut remodelling in response to mating and diet and very little lifespan response to DR (Magwere *et al*., 2004).

The gut is not only an important signalling centre, which may influence the rate of aging in other cells and tissues, but it is also an important barrier to infection as well as the tissue which mediates all nutrient absorption into the body (Rera *et al*., 2012; Miguel-Aliaga *et al*., 2018). Moreover, the appearance of age-associated gut dysfunction is closely correlated with death in females (Rera *et al*., 2012). The exact process through which age-related gut dysfunction occurs is not fully understood, however, it is thought that over the course of a lifetime, intestinal stem cell (ISC) division slowly becomes dysregulated, leading to hyperplastic intestinal pathologies. This over-proliferation of ISCs leads to a build-up of undifferentiated enteroblasts – the product of this is cell crowding and eventual tumour formation (Choi *et al*., 2008; Biteau *et al*., 2010; Patel *et al*., 2015). This can inhibit sufficient nutrient absorption and lead to an eventual loss of intestinal barrier function, exposing the flies to malnutrition and possible lethal infections (Choi *et al*., 2008; Biteau *et al*., 2010; Patel *et al*., 2015).

Interestingly, age-associated gut pathologies are responsive to diet (Choi *et al*., 2008; O’Brien *et al*., 2011; Regan *et al*., 2016), suggesting that DR extends lifespan by reducing gut pathology and so preserving late life gut health. Lifespan extension under DR is often associated with a reduction in signalling from the nutrient sensitive, molecular growth pathway, Target of Rapamycin (mTOR) (Fan *et al*., 2015). Treatment with the mTOR inhibitor Rapamycin increases intestinal barrier function during aging, and extends lifespan (Schinaman *et al*., 2019; Yu *et al*., 2025). Further, gut pathologies are ameliorated by DR and/or administration of Rapamycin (Regan *et al*., 2016; Regan *et al*., 2022; Yu *et al*., 2025). It has been proposed that inhibiting mTOR signalling slows down the proliferation rate of ISCs in ageing guts (Fan *et al*., 2015). In this model, DR extends lifespan as a function of mTOR signalling inhibition, which prevents ISC over-proliferation and sustains gut health for longer (Fan *et al*., 2015).

Our recent work has established that an essential dietary micronutrient, a sterol, is the key determinant of lifespan in response to DR in *Drosophila* females (Zanco *et al*., 2021). These data showed that lifespan is longer on lower protein (or DR) diets because flies on DR food are rescued from a sterol deficiency, which may prevent somatic damage. In fact, even flies fed a high protein diet, which both increases TOR activity and shortens lifespan, have longer lives when the diet is supplemented with sterols, and their longevity is almost equivalent to that caused by DR or rapamycin supplementation (Zanco *et al*., 2021). Thus, rapamycin supplementation may be beneficial primarily because it reduces growth and/or reproduction, which then indirectly rescues flies from a life-shortening micronutrient deficiency. Thus, if gut integrity is key to longer life, then dietary sterols should be important for the occurrence of later life gut pathologies in *D. melanogaster*.

In the current study we investigated the hypothesis that sterol-depleted diets target intestinal health as a precursor to altering lifespan. To do this we varied the concentration of sterols in the diet of female *D*.*melanogaster*. We then assessed lifespan, gut function using Smurf assays, and examined epithelial cellular disruptions through microscopy.

## Results

### Low dietary sterol intake is associated with increased intestinal permeability and disruptions to the midgut epithelium

Preventing sterol depletion in flies may mediate the lifespan benefits of DR by preserving age-related intestinal integrity. To assess this, we fed flies a diet containing the food dye FD&C Blue Dye number 1 throughout adult life, which by itself had no effect on lifespan (Figure 1a, Supplementary Table 2) and allowed us to directly measure if and when gut integrity is lost by measuring the leakage of dye from the gut into the body cavity (“smurfing”)(Martins *et al*., 2018). Flies were considered smurfs if their whole body turned completely blue, as described by Martins *et al (2018)*, ‘light smurfs’, which are only partially blue were not recorded as smurfs.

**Figure 1.**
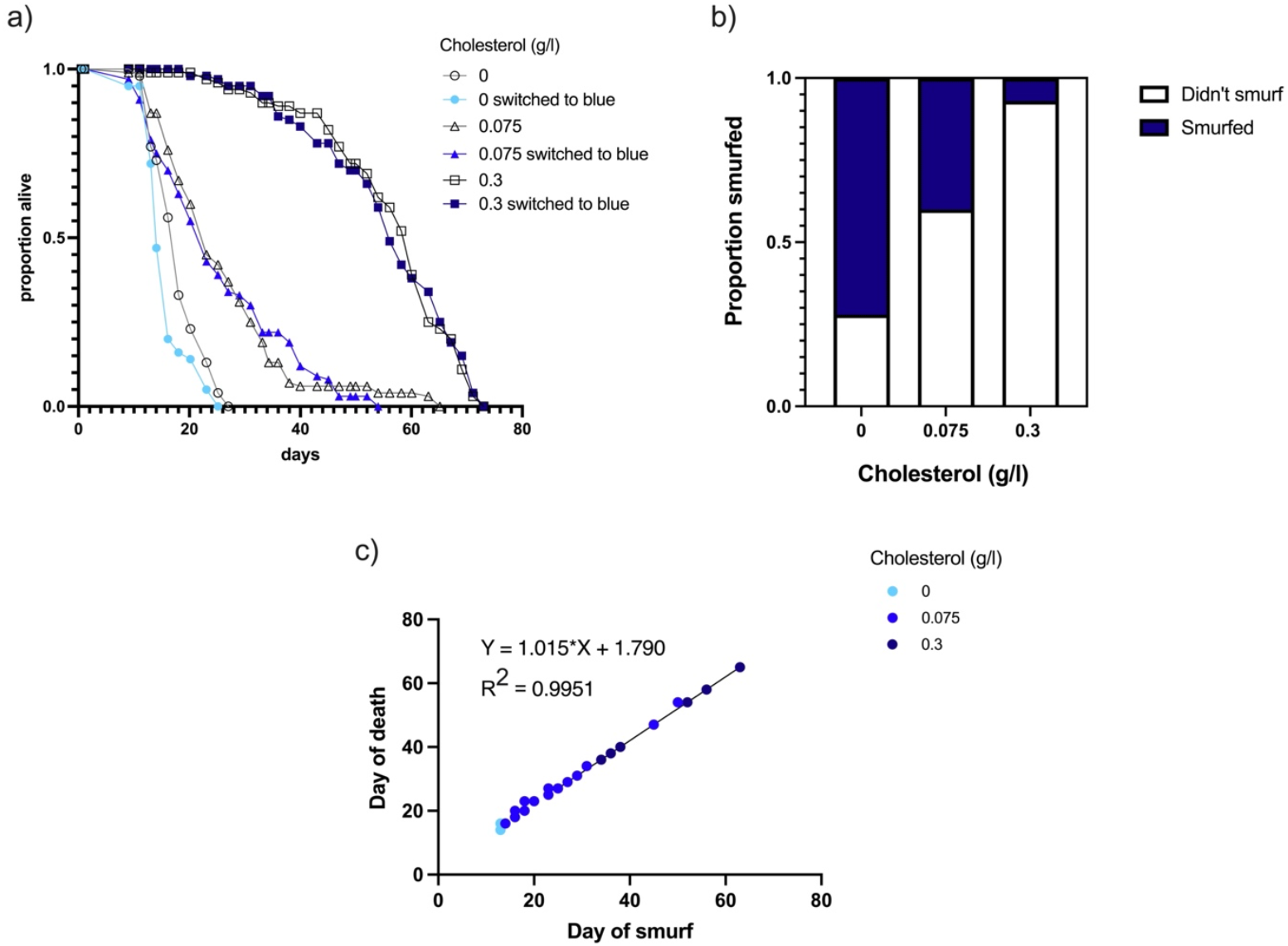
Flies fed low sterol diets have greater levels of gut permeability than those fed high sterol diets. Lifespan was maximised on a high cholesterol diet and declined as the cholesterol concentration was reduced (a). For each cholesterol concentration, one cohort of flies were either switched to blue food on day ~ 12 from adult emergence or kept on the same diet without blue dye (b). Blue dye did not have a significant effect on lifespan (p = 0.060). Cholesterol has a significant effect on smurfing in flies prior to death (p = <0.001) (b). This increase in smurfing is negatively correlated with the cholesterol concentration of the diet (b). In flies that do smurf, death occurs within ~ 48hrs of turning blue, irrespective of dietary cholesterol intake (c). All statistics are reported in Supplementary Tables 1 and 2. To assess intestinal permeability, we fed flies a diet containing the food dye FD&C Blue Dye number 1 throughout adult life. This allowed us to directly measure if and when gut integrity is lost by measuring the leakage of dye from the gut to permeate throughout the body (“smurfing”)(Martins et al., 2018). Flies were considered smurfs if their whole body turned blue.

We found that both lifespan decreased and the proportion of flies that smurfed before dying increased as sterols were diluted in the diet (Figure 1a,b, Supplementary Table 1 and 2). Specifically, 72% of flies on diets containing 0g/l of dietary cholesterol (Figure 1b) smurfed before dying, with a population median lifespan of 15d. On diets containing 0.075 g/l cholesterol, 40% of flies smurfed before dying, with population median lifespan of 24d. While only 7% of flies with 0.3g/l of cholesterol in the food smurfed before dying, with a population median lifespan of 57d (Figure 1a). Interestingly if flies did smurf, the onset of gut permeability consistently occurred ~48hrs before death (Figure 1c) irrespective of the dietary sterol level, indicating a threshold effect, such that when the gut became permeable to the blue dye, death was inevitable and imminent. These findings suggest that loss of intestinal barrier function contributes to the higher death rate caused by dietary sterol depletion, but they also show that not all flies develop this form of gut permeability before dying.

### A low sterol diet is associated with increased disruptions to the anterior midgut epithelium when compared to diets containing adequate sterols

Similar to the mammalian digestive tract, the midgut of adult *Drosophila* is highly compartmentalised and can be divided into regions with distinct morphological, histological, and genetic properties (Veenstra, Agricola and Sellami, 2008; Buchon *et al*., 2013; Dutta *et al*., 2015). Of these regions, we know that both the cellular structure and arrangement of epithelial cells in these regions are modified by diet (Regan, *et al*., 2016). Because of this, we examined the effects of sterol limitation on both the anterior and posterior regions of the midgut in flies that were 18 days old (Figure 2, Supplementary Table 3). This time point was chosen as it aligns with the commencement of death and smurfing on low sterol diets (Figure 1b,c).

**Figure 2.**
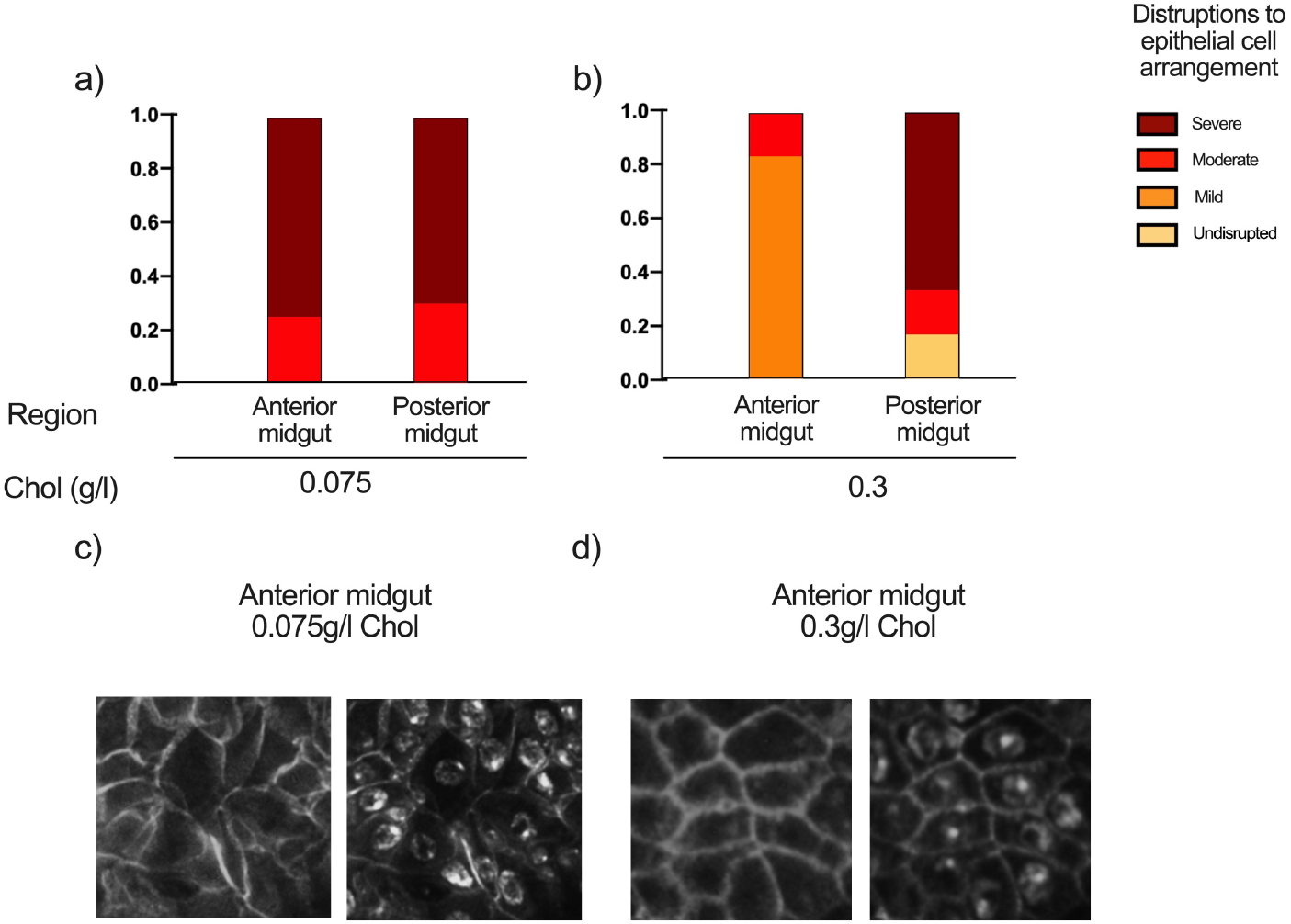
Disruptions in the anterior midgut epithelium are sterol dependent (a,b). Disruptions in the epithelial cells of the Drosophila midgut at 18 days of age were significantly greater in flies fed low sterol diets (a,c, Supplementary Table 3-4) when compared with flies that were fed an adequate supply of sterols (b,d, Supplementary Table 3-4). In contrast, the posterior midgut did not present with significantly different classifications of epithelial disruption across both sterol treatments (a, b, Supplementary Table 3-4). Epithelial cell images were de-identified and binned into scaled categories based on visual appearance of cell organisation, where 1 = undisrupted honeycomb cells arranged with well aligned nuclei, 2 = a mixture of honeycomb structured cells and early-stage disorganisation, <50% of cells are elongated and irregular shaped, 3 = highly disorganized with > 50% of cells are elongated and irregular shaped, and 4 = severe disruptions in epithelial cell organisation, all cells are misaligned, elongated and irregular in shape. These categories were adapted from a scoring system for epithelial disorganisation published by Regan et al. (2016).

Guts were stained with phalloidin and DAPI and imaged using a CV1000 confocal microscope, so cellular organisation could be scored. All images were de-identified and binned into scaled categories based on visual appearance of the epithelium (described in methods). Healthy epithelial cells have a honeycomb arrangement with perfectly aligned nuclei, while unhealthy epithelial cells present as misaligned and elongated or irregular in shape (Regan, *et al*., 2016). We found that flies fed low sterol diets showed significantly more signs of epithelial cell disruptions in the anterior midgut (Figure 2a, Supplementary Table 3), while flies fed adequate sterols had well-maintained cells in this region (Figure 2b, Supplementary Table 3). Interestingly, there was no significant difference in cellular organisation in the posterior midgut, with flies fed both low and adequate sterols presenting with signs of severe pathologies in this region (Figure 2a,b, Supplementary Table 3 - 4). The anterior midgut data therefore associate closely with the Smurf assay results, showing a relationship with early versus late mortality. In contrast, posterior midgut disorganisation appears to occur at an equal pace between treatments regardless of future life expectancy, suggesting that the types of structural changes in epithelial cells scored in the posterior midgut are unlikely to be limiting factors for longevity.

### Antibiotics do not modify the lifespan limiting effects of a low sterol diet

Reduced gut barrier function may cause death by allowing bacteria to infiltrate the body cavity, causing sepsis. We checked to see if antibiotic treatment could thus increase the lifespan of flies fed low sterol diets. Treatment with a cocktail of five antibiotics that have been shown to clear bacteria that normally associate with *D. melanogaster* larvae (Consuegra *et al*., 2020a; Consuegra et al., 2020b) had a small, but significant positive effect on lifespan overall, but did not modify the way that sterol restriction shortened lifespan (Figure 3, Supplementary Table 5). This indicates that the more rapid death upon sterol depletion is not due to sepsis caused by these bacteria, and therefore may indicate that gut barrier function either causes death by other means (e.g. loss of absorptive capacity) or gut permeability is a symptom, rather than a cause, of the progressive increase in mortality in these flies.

**Figure 3.**
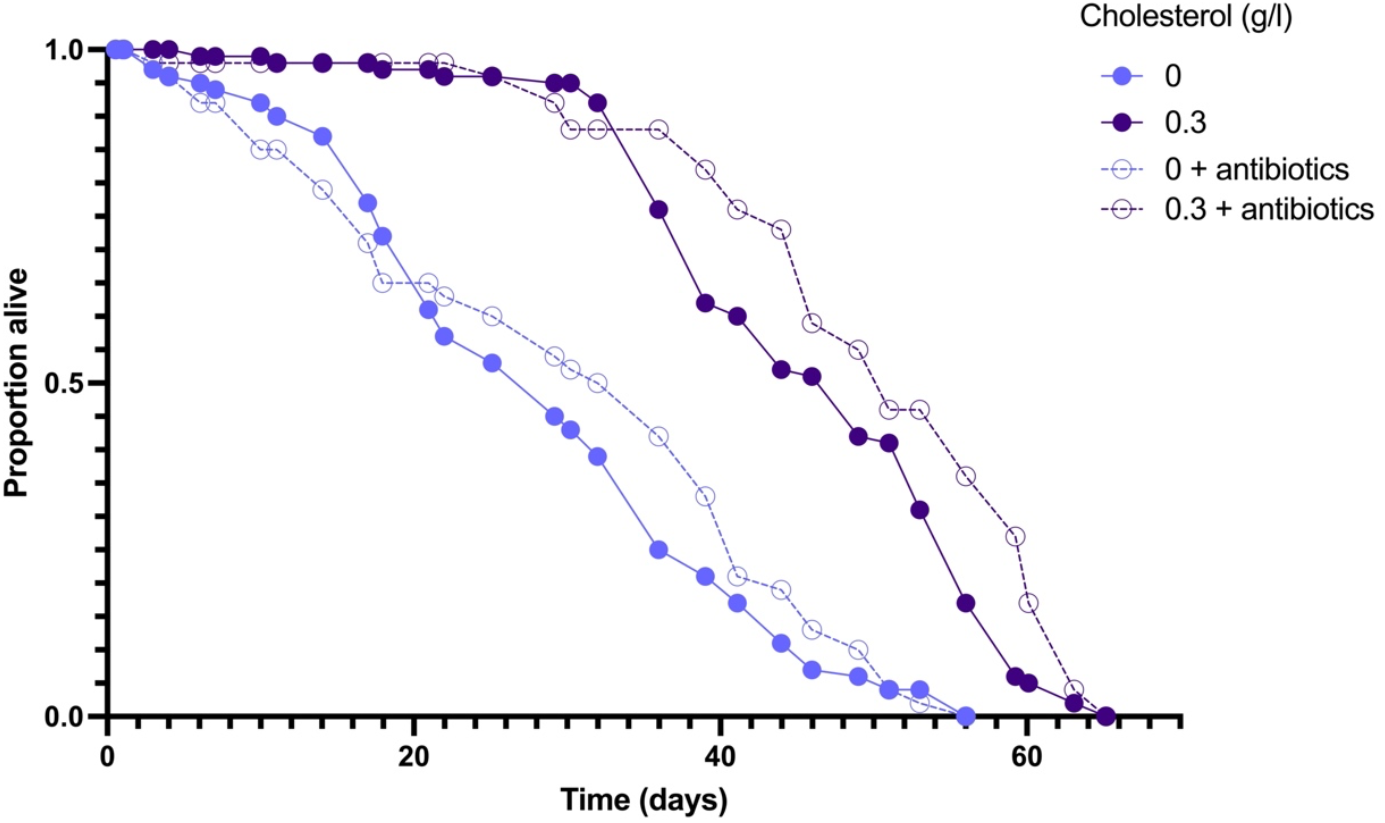
Antibiotic treatment in sterol-limited flies does not rescue lifespan. Flies were fed throughout life on either a sterol sufficient or sterol deficient diet, with or without antibiotics. Lifespan data for each treatment was analysed using a cox proportional hazards model. There was a significant effect of both antibiotics (p = 0.003) and cholesterol (p = <0.001) on lifespan, but no interactive effect (p = 0.872). All statistics are reported in Supplementary Table 5.

## Discussion

Trade-offs are a fundamental tenet of evolutionary life history theory. The lifespan benefits of DR regimes that reduce reproduction has led to the conclusion that reproduction is locked into an inevitable trade off against somatic maintenance, which underpins ageing, due to competition for the same limiting resource (Maynard Smith, 1958). This concept is the foundation of the resource reallocation theory of ageing (Kirkwood and Holliday, 1979; Holliday, 1989; Kirkwood and Rose, 1991; Hughes and Reynolds, 2005). Despite the long history of this theoretical description, the molecular basis of this trade-off between lifespan and reproduction has remained elusive.

We now know that limiting dietary sterol levels can account for the lifespan costs observed for *Drosophila* feeding on high yeast (protein) diets, thus explaining why restricting the diet can provide a comparative benefit at least in this species (Zanco *et al*., 2021). We also know that in female flies, the ovaries require a constant supply of sterols for egg production and this supply can come from the diet and/or internal reserves (Heier *et al*., 2021). Thus, when sterol deprived, it is possible that they initiate a program of sterol mobilisation from somatic tissues, which includes the gut, to ensure maintenance of a high level of reproductive output (Zanco *et al*., 2022). If sufficiently severe, dietary sterol depletion may cause lethal damage to the mother if she retrieves sterols from the gut to the extent that barrier and absorptive function is lost (Piper *et al*., 2022). Thus, DR may preserve lifespan by rescuing mothers from damage caused to the gut by sterol depletion.

In the current study we investigated the hypothesis that sterol-depleted diets target intestinal health as a precursor to altering lifespan. In support of this model, we found that gut integrity does indeed decline at an increasingly higher frequency as the dietary protein to sterol ratio increases, and this is associated with progressively shortened lifespan. Although we did not examine incidence of smurfing in males, we know that sterol depletion does not limit male lifespan (Wu *et al*., 2020). We also know that males do not smurf when fed a high protein diet (Regan *et al*., 2016), thus, it can be assumed that our findings represent a sexually dimorphic effect of nutrition on lifespan and reproduction in *D. melanogaster* (Magwere *et al*., 2004; Regan *et al*., 2016; Wu *et al*., 2020).

While our data also show a strong connection between gut function and death during DR, they also demonstrate that this link is specific to conditions of sterol depletion, and suggest that although loss of intestinal barrier function is associated with ageing in numerous organisms from worms to primates (Martins *et al*., 2018; Salazar *et al*., 2023), it is not a universal explanation for age-related death. Indeed, the majority of our flies that died of old age under fully fed conditions did not smurf before dying, which is in line with findings by others that have uncoupled lifespan and gut function (e.g. Rera *et al*., 2012; Fan *et al*., 2015; Regan *et al*., 2016; Wu *et al*., 2016; Schinaman *et al*., 2019). Thus, although we observe gut dysfunction in broad range of taxa in old age, it is unlikely to be a conserved physiological cause of death.

The model we have proposed identifies sterols as a limiting currency that is exchanged in the lifespan/reproduction trade-off under DR in *Drosophila*. Interestingly, while we were preparing this manuscript, another study has also shown that dietary sterol limitation increases gut permeability and shortens lifespan, and that these effects are suppressed by genetic ablation of egg production or administration of rapamycin (Yu *et al*., 2025). Flies, and other insects, may be particularly prone to this nutrient limitation as they are sterol autotrophs (Behmer and Nes, 2003). Moreover, their natural source of sterols is yeast, which fluctuate in sterol composition depending on the environment in which they grow (Becher *et al*., 2012; Markow *et al*., 2015). This situation is even more pronounced by the marked variation in diet composition between labs in which yeast can be supplied in forms that vary from water soluble yeast extracts (high in protein, but sterol depleted) through to yeast products containing whole-cells and their lysates (high in protein and containing sterols) (Bass *et al*., 2007; Lesperance and Broderick, 2020; Zanco *et al*., 2021). Thus, fly diets in which protein concentration is elevated by increasing yeast extract levels would create a greater imbalance between protein and sterol supply compared to diets using whole cell lysates. Supporting this observation, the effects of DR on lifespan tend to be more pronounced when yeast extract is used rather than whole cell lysates (Bass *et al*., 2007; Piper and Partridge, 2007).

Many organisms that exhibit lifespan responses to DR are sterol heterotrophs, meaning they can synthesise their own sterols. Thus, sterol limitation is unlikely to mediate the effects of DR across different taxa. However, our data may point to a general principle that when organisms commit to reproduction based on a limited subset of nutrients (e.g. just the macronutrients (Simpson and Raubenheimer, 2012; Solon-Biet *et al*., 2014)), they are at risk of suffering from critical shortages in other essential nutrients that they then deplete in a way that decreases lifespan. In turn, we might expect that different animals are pre-disposed to different micronutrient deficiencies. For example, there is evidence that rodents, and perhaps humans, remobilize calcium from their bones and teeth to supplement infant nutrition during lactation when dietary calcium intake is low (Picciano, 2003; Prentice, 2003; Miller and Bowman, 2004; Speakman, 2008), and therefore the molecular processes underlying these trade-offs are likely to be organism specific.

## Materials and Methods

### Fly husbandry and strains

All flies were reared and mated prior to experiments following the protocol described in Zanco et al. (2021). Mated females were then placed on holidic diets varying in sterol concentrations – recipes described in Zanco et al. (2021) for the duration of their lifespans. A *w*^*1118*^ strain (Cat#: 60000) from Vienna Drosophila Resource Center (VDRC, Vienna, AUT) was used to examine the effects of antibiotics on lifespan. All other experiments were conducted using a wild type *Drosophila melanogaster* strain called Dahomey (Mair, Piper and Partridge, 2005).These flies have been maintained in large numbers with overlapping generations to maintain genetic diversity. All flies were reared for two generations at a controlled density before use in experiments, to control for possible parental effects. Eggs for age-synchronised flies were collected over 18 hr, and the resulting adult flies emerged during a 12 hr window. They were then allowed to mate for 48 hr before being anaesthetised with CO_2_, at which point females were separated and allocated into experimental vials. Stocks were maintained and experiments were conducted at 25 °C on a 12 hr: 12 hr light:dark cycle at 65% humidity (Bass *et al*., 2007).

### Lifespan assays

For all lifespan assays, flies were placed into vials (FS32, Pathtech) containing 3 ml of treatment food at a density of ten flies per vial, with ten replicate vials per treatment. Flies were transferred to fresh vials every two to three days at which point deaths and censors were recorded and saved using the software package Dlife (Linford *et al*., 2013; Piper and Partridge, 2016).

### Intestinal barrier function analyses

The protocol described in Martins *et al*. (2018) was adapted as a means of assessing intestinal barrier function. On ~ day 12 from adult emergence, flies from each dietary treatment group were switched to experimental diets of the same recipe but with the addition of blue food dye (FD&C blue #1). Flies were then checked every second day for “smurfing”. This phenotype is caused by the blue dye escaping the gut lumen and entering the body cavity, which turns the whole fly blue. Flies were recorded as smurfs when the entire fly was blue.

### Immunohistochemistry

On day 18 (when flies had begun to die) guts were dissected from live flies (at least four per treatment) in ice-cold PBS and immediately fixed in 4% paraformaldehyde (sigma) for 60 minutes at room temperature. After fixing, guts were washed with 0.1% Tween (PBST) then blocked for 30 min at room temperature in 2% heat-inactivated normal donkey serum in PBT. Guts were stained with DAPI at a concentration of 5 uL/mL and Phalloidin at a concentration of 14uL/mL. All guts were mounted in Fluoromount-G (SouthernBiotech). Regions of interest in the midgut were then imaged using a confocal microscope (Olympus CV1000) with a 40x objective lens.

A minimum of 4 guts were imaged for each region of the gut 1 and 2 (anterior) and 4 and 5 (posterior), per condition. Guts were excluded from analyses if specimen or image quality were poor. An area of 150µm * 150µm (centered) was selected using Fiji, and then cells in this region were binned into scaled categories, where 1 = undisrupted honeycomb cells arranged with well aligned nuclei, 2 = a mixture of honeycomb structured cells and early-stage disorganisation, <50% of cells are elongated and irregular shaped, 3 = highly disorganized with > 50% of cells are elongated and irregular shaped, and 4 = severe disruptions in epithelial cell organisation, all cells are misaligned, elongated and irregular in shape. These categories are adapted from a scoring system for epithelial disorganisation published by Regan et al. (2016). These regions were selected as they are known to be impacted by dietary restriction (Rera et al. 2016). All images were de-identified before scoring.

### Antibiotic assay

To examine the role of sepsis in death following loss of gut barrier function, antibiotics were added to the fly food. Mated females were placed on a holidic medium varying in cholesterol concentrations and antibiotic presence (Tetracylin, Ampicillin, Kanamycin, Erythromycin) for the duration of their lifespans. The antibiotic mix was prepared as described by (Consuegra *et al*., 2020). Each antibiotic was added at a concentration of; kanamycin (50 μg/mL), ampicillin (50 μg/mL), tetracycline (10 μg/mL) and erythromycin (5 μg/mL). Stock solutions of each antibiotic were made in either MilliQ water (ampicillin and kanamycin) or ethanol (erythromycin and tetracycline) based on their solubility, such that adding 1mL of stock to 1L holidic media resulted in the above final concentrations. Immediately before dispensing the media, each antibiotic solution was added with a micropipette, under constant stirring to promote even mixing. A separate length of tubing was used to dispense the food containing antibiotics, ensuring no antibiotics were inadvertently introduced to the antibiotic free media.

### Statistical analyses

All statistical analyses were performed using R Studio (Version 1.2.5042), available from http://www.R-project.org/). For lifespan experiments, the median lifespan was obtained for each experimental vial prior to analysis. All data were analysed using Generalised Linear Mixed Effects Models. Posthoc tests using emmeans were then applied to models for epithelial cell and smurf data.

