## Supplementary Table 1. for "Dietary sterol depletion is associated with gut dysfunction in *Drosophila melanogaster* females"

Supplementary Table 1. *A generalized linear model with replicate vial used as a random effect, reveals that cholesterol has a significant effect on smurfing.*

| Smurfing | Chisq | DF | Pr(>Chisq) |
|---|---|---|---|
| Cholesterol | 21.976 | 1 | <0.001*** |
