## Supplementary Table 2. for "Dietary sterol depletion is associated with gut dysfunction in *Drosophila melanogaster* females"

Supplementary Table 2. *The effects of cholesterol and blue dye on median lifespan were analysed using a linear mixed effects model, with replicate vial used as a random effect. There was a significant effect of cholesterol on lifespan, but no significant effect of blue dye, nor an interactive effect between cholesterol and blue dye.*

| Median lifespan | Chisq | Df | Ps(>Chisq) |
|---|---|---|---|
| Cholesterol | 345.99 | 2 | <0.001*** |
| Blue dye | 3.65 | 1 | 0.060 |
| Chol : Blue dye | 3.45 | 2 | 0.177 |
