## Supplementary Table 3. for "Dietary sterol depletion is associated with gut dysfunction in *Drosophila melanogaster* females"

Supplementary Table 3*. The effects of cholesterol on epithelial cell organisation in the midgut were analysed using a linear mixed-effects model, where gut was used as a random effect. There was a significant effect of both dietary cholesterol and the interactive effect between cholesterol and gut region on epithelial cell disturbance.*

| *Midgut epithelial cell disturbances* | Chisq | Df | Pr(>Chisq) |
|---|---|---|---|
| Cholesterol | 10.732 | 1 | <0.001*** |
| Region | 0.697 | 1 | 0.404 |
| Cholesterol : Region | 4.765 | 1 | 0.030* |
