## Supplementary Table 4. for "Dietary sterol depletion is associated with gut dysfunction in *Drosophila melanogaster* females"

Supplementary Table *4. The effects of cholesterol in the anterior and posterior regions of the midgut were analysed using a Posthoc estimated marginal means (emmean) test on the linear-mixed effects model above (Table 3). The effect of cholesterol on epithelial cell disturbance was significantly different to all other conditions in the anterior guts of flies fed 0.3g/l cholesterol.*

| *Region* | Cholesterol (g/l) | emmean | SE | Df | lower.CL | Upper.CL | Group |
|---|---|---|---|---|---|---|---|
| Anterior | 0.3 | 2.20 | 0.318 | 18.0 | 1.53 | 2.86 | 1 |
| Anterior | 0.3 | 3.42 | 0.282 | 20.5 | 2.83 | 4.01 | 2 |
| Posterior | 0.075 | 3.66 | 0.252 | 17.4 | 3.13 | 4.19 | 2 |
| Posterior | 0.075 | 3.72 | 0.397 | 19.9 | 2.89 | 4.55 | 2 |
