## Supplementary Table 5. for "Dietary sterol depletion is associated with gut dysfunction in *Drosophila melanogaster* females"

Supplementary Table 5. *The effects of antibiotic administration on median lifespan were analysed using a linear mixed-effects model, with replicate vial used as a random effect. There was a significant effect of both antibiotic administration and cholesterol on lifespan, but no significant interactive effect between antibiotics and cholesterol.*

| Median lifespan | Chisq | Df | Pr(>Chisq) |
|---|---|---|---|
| Antibiotics | 8.907 | 1 | 0.003** |
| Cholesterol | 190.511 | 1 | <0.001*** |
| Cholesterol : Antibiotics | 0.026 | 1 | 0.872 |
